# High-purity stem cell-derived β-cells recapitulate key transcriptional and functional features of human islets

**DOI:** 10.64898/2026.05.22.726825

**Authors:** Rémi B. Fiancette, Junyue Huang, Christine Stephens, Jessica E. Hibbert, George Hewitt, Christopher Carlein, Ali H. Shilleh, Charlotte Clinton, Luana De Abreu Queiros Osorio, David S. Tourigny, Steven J. Millership, Victoria Salem, David J. Hodson, İldem Akerman

## Abstract

Human pluripotent stem cell-derived islets (SC-islets) offer an excellent medium for human pancreatic disease modelling and mechanistic studies into diabetes. While substantial progress has been made in differentiation protocols, their implementation in different laboratories result in variable β-cell proportions with contaminant non-endocrine and proliferative cell types. To date, no facility-level implementation exists for producing SC-islets that can be shipped and benchmarked across multiple sites. Here, we describe the scalable optimisation, standardization, and facility-level implementation of an established human stem cell differentiation strategy that consistently results in a high proportion of β-cells, with up to 75% of cells co-expressing C-peptide and the pancreatic endocrine marker, ISL1. Functionally, SC-islets exhibit glucose-responsive calcium influx and insulin secretion, recapitulating key physiological β-cell functions. Single-cell transcriptomic profiling reveals a simplified endocrine landscape dominated by β-cells, with a striking transcriptional similarity to human primary β-cells (Pearson’s *r*^2^∼0.9). We observe smaller fractions of α- and enterochromaffin-like cells with very low levels of poly-hormonal or proliferating cell types (<3%). Taken together, we provide a well-defined, reproducible and accessible *in vitro* SC-islet platform benchmarked for functionality at multiple recipient sites.

## INTRODUCTION

Diabetes mellitus results from a failure to secrete enough of the hormone insulin which is made and secreted by the β-cells located in the pancreatic islets of Langerhans. Diabetes is a major health concern that affects >400 million people worldwide^1^, thus β-cell biology is a key focus of biomedical research. In recent years, significant progress has been made in the development of protocols^2–7^ that can transform pluripotent stem cells into pancreatic islets (SC-islets), offering an excellent *in vitro* model for the investigation of human-specific regulatory mechanisms of pancreatic development and endocrine pancreas disorders^8–16^. These cells also hold promise for cell replacement therapies for type 1 diabetes (T1D)^17–20^, with potential benefits extending to type 2 diabetes^21^ treatments. Despite these advances, research utilising SC-islets remains largely confined to sufficiently equipped laboratories, as non-specialised groups face significant challenges in generating quality-controlled cells due to the infrastructure and expertise required. Therefore, access to a standardized, consistent and a reliable source of functional SC-islets would facilitate cell replacement research for T1D, provide a non-animal model for studying β-cell function and endocrine pancreas development particularly for researchers without stem-cell biology expertise.

Protocols for differentiating human pluripotent stem cells into glucose-responsive, insulin-secreting β-like cells typically generate cultures containing ∼35-50% β-cells, with most recent protocols reaching >60% (Table S1). In depth analysis of SC-islets reveals that they remain heterogeneous, with laboratory-dependent β-cell yields that contain varying proportions of non-β cell populations, including mesenchymal, epithelial, ductal, neural crest–like cells as well as proliferating cells^7,22–29^. Numerous strategies have been developed to enrich β-cells from stem cell–derived islet preparations, including approaches based on surface marker selection and small-molecule treatments^30–38^. While these methods can improve endocrine purity and function, they come at the cost of reduced yield, increased expense, and potential impacts on cellular viability. This highlights the need for cost-effective differentiation platforms that intrinsically generate high proportions of β-cells without requiring downstream enrichment steps.

Here, we describe the optimization and standardization of a published^7^ SC-islet differentiation strategy using small scale bioreactors within a dedicated academic research facility. We demonstrate consistent and high β-cell yields across lines and batches, with transcriptomic and functional assays reliably supporting SC-islets as a valuable model for research. Notably, our approach focuses on high reproducibility and β-cell content with only low or absent non-endocrine contaminant or proliferating cell populations. Single cell transcriptomic analysis shows a profound similarity between the transcriptional landscape of SC-islet and primary islet cell types, while also highlighting pathways for future refinements of the protocol.

## RESULTS

### Reproducible β-cell generation from stem cells

Access to consistently produced SC-islets for researchers lacking the necessary expertise or facility remains a key challenge in the field. To enable reproducible production of SC-islets, we standardized upstream human embryonic stem cell (hESC) maintenance, enforcing a long duration, fixed-interval feeding regimen aimed at minimizing variability before lineage induction. Following this standardized expansion phase, hESC H1 cells were differentiated using a modified multi-stage protocol^7^ in 30 mL magnetic stir-based bioreactors (Figure 1A, Methods), and differentiation efficiency was assessed at key developmental checkpoints by flow cytometry using well-established markers for each stage. In the hESC H1 line, definitive endoderm formation is highly efficient, with an average of 93% of cells co-expressing the endodermal markers FOXA2 and SOX17 (day3; Figure 1B). Cells complete the pancreatic progenitor stage with an average of 80% double positive for the lineage-defining markers PDX1 and NKX6-1 (day12; Figure 1C). At the final maturation stage, we observe a substantial enrichment of β-cells with an average of 68% of cells double positive for the endocrine and β-cell markers Islet-1 (ISL1) and C-peptide (day37, Stage7w2; Figure 1D). In the highest-performing differentiations, this proportion reaches >75%, representing a significant increase in β-cell proportion compared with published outcomes using the original protocol (Table S1). SC-β cells produced by this protocol are also ∼60% double positive for C-peptide and NKX6-1, a commonly used β-cell marker (Figures 1D and S1A). Preferential differentiation toward the β-cell lineage in our preparations comes at the expense of contaminant cells as well as pancreatic α-cells, with only ∼5% of cells co-expressing ISL1 and the peptide hormone glucagon (GCG) (Figures 1D and S1A). Importantly, glucagon was restricted to the ISL1^+^ (endocrine) cells with < 3% C-peptide and glucagon double-positive cells as assessed by flow cytometry (Figure S1A). This suggests that bihormonal cells are negligible at the protein level, and therefore unlikely to confound functional assays.

**Figure 1.**
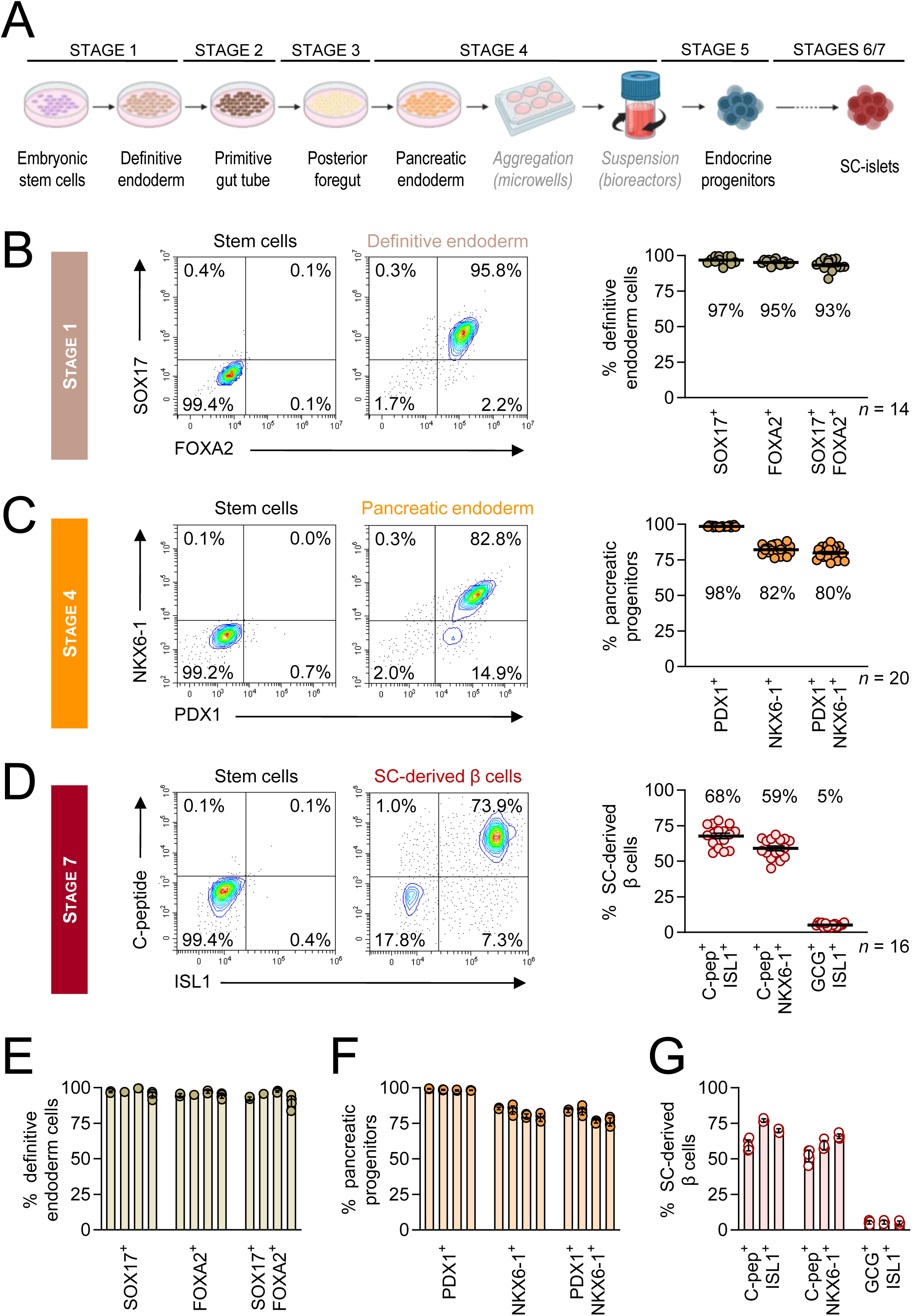
Standardized differentiation of hESCs yields reproducible β-cell enrichment. (A) Schematic of the 7-stage SC-islet differentiation protocol (created with BioRender.com). (B-D) Representative flow cytometry contour plots (left panels), and quantification across independent bioreactors (right panels), showing marker expression at (B) definitive endoderm (Day3, FOXA2/SOX17; n = 14), (C) pancreatic progenitor (Day12, PDX1/NKX6-1; n = 20), and (D) maturing SC-islet stage (Stage7 week2, ISL1/C-peptide; NKX6-1/C-peptide; ISL1/Glucagon; n = 16) derived from hESC1 (H1) cell line. (E-G) Quantification of marker expression at key differentiation stages across different batches of production (E) endoderm markers, (F) pancreatic progenitor markers, and (G) mature SC-β cell markers. Each dot represents an individual bioreactor, while bars in (E-G) denote individual differentiation batches, illustrating intra-and inter-batch consistency. Error bars denote mean ± SEM with the exact values annotated in (B-D) and mean ± SD in (E-G).

To assess the consistency of β-cell generation, we quantified variability in the proportion of C-peptide positive cells across independent differentiation runs and between bioreactors within each run (Table S2, Figures 1E-1G). While some batch-to-batch variation is observed, intra-batch variability remains low, typically within ∼5%, indicating that cells differentiated in parallel can be compared with confidence and the platform provides sufficient reproducibility for genetic perturbation and disease modelling studies.

Comparable differentiation efficiencies are obtained using an independent hESC line (hESC3, H3 where a Green Fluorescence Protein reporter gene has been targeted into one of the alleles of the *INS* locus^39^), yielding similar proportions of endodermal (94%), pancreatic progenitor (82%), and SC-islet populations (46% β- and 14% α-cell, Figures S1B-S1E), supporting the robustness of this standardized approach across genetic backgrounds.

### Stem cell–derived β-cells exhibit glucose responsiveness and stimulus–secretion coupling

We next evaluated the properties of our SC-islets to assess their suitability as a model of human β-cell function. On average, SC-islets derived from the H1 line yield ∼5000 clusters, corresponding to ∼18 million cells per 30 mL bioreactor, representing an overall yield of ∼1.4-fold relative to the starting cell numbers (Figures 2A-2B and S2A). As expected, H3-derived SC-islets start to express reporter GFP from the endocrine progenitor stage, concomitantly with the induction of insulin expression (Figure S2A). Total insulin protein content was similar between SC-islets derived from the H1 and H3 lines (Figure 2C) and was comparable to donor-derived islets (Figure S2B). This is consistent with scRNA-sequencing data showing reduced but broadly comparable insulin mRNA levels in stem cell-derived (SC-) and primary β-cells (Figure 2D).

**Figure 2.**
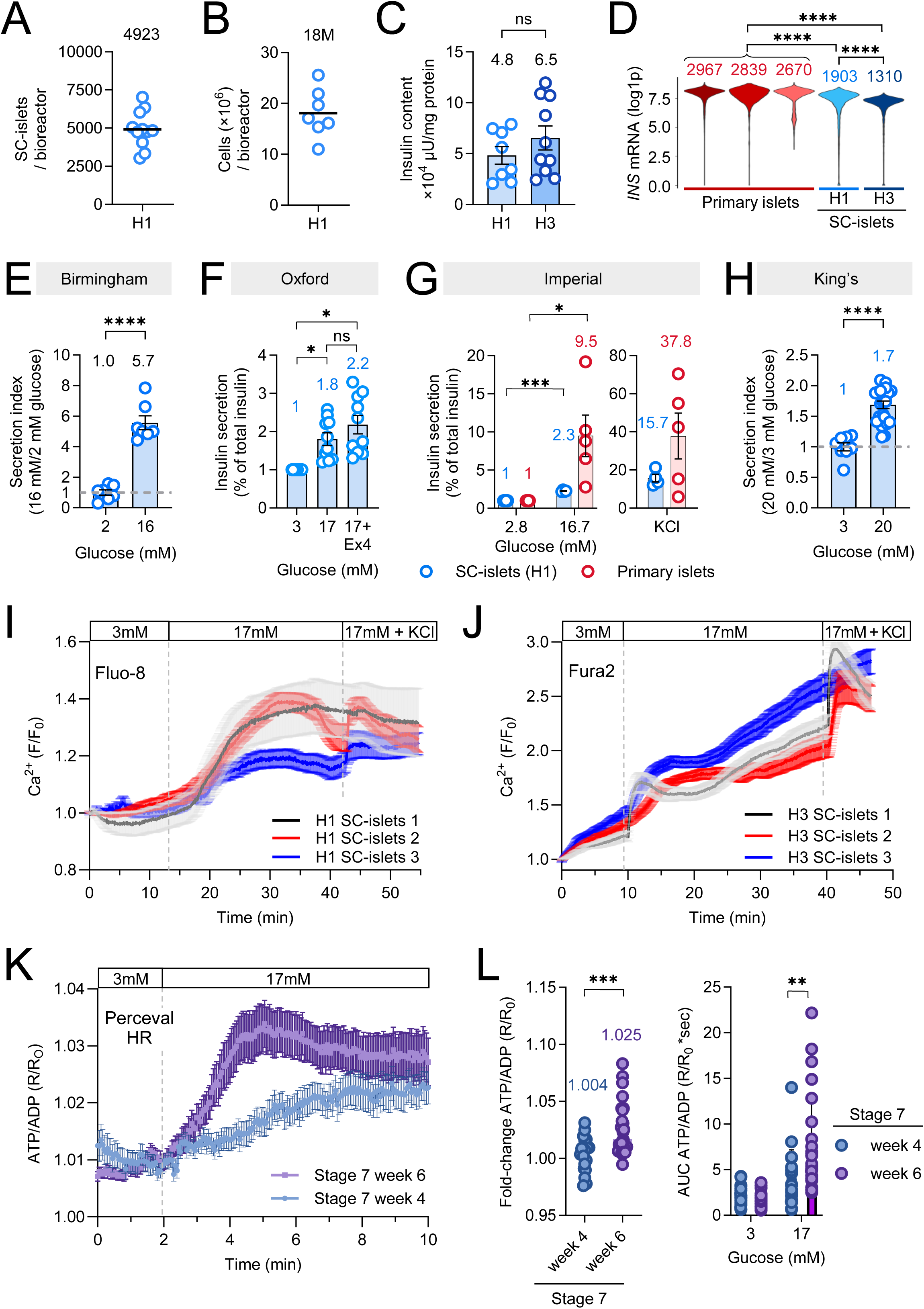
SC-islets recapitulate key β-cell functions. (A) Number of SC-islet clusters per 30 mL bioreactor (n = 11 bioreactors from 2 differentiation batches) (B) Number of cells per bioreactor (n = 7 bioreactors from 2 differentiation batches) derived from H1 hESC line. (C) Quantification of insulin content in H1 and H3 hESC-derived SC-islets (Stage7w2, n = 8 and 10 bioreactors, respectively). (D) Violin plots showing INS expression within three donor-derived human islet samples (primary islets) and SC-islets derived from H1 and H3 cell lines (log-normalized RNA counts). Annotations are linear normalized RNA counts. (E) Static glucose-stimulated insulin secretion (GSIS) assay performed at the production site (Birmingham), demonstrating glucose-responsive insulin release. Insulin secretion is assessed in SC-islets at Stage7w2 by ELISA and normalized to total insulin content and expressed relative to basal conditions (2 mM glucose vs 16 mM glucose in the presence of IBMX; n = 7 bioreactors from 3 differentiations). (F-H) Functional characterization of H1 hESC-derived Stage 7 SC-islets by GSIS assay and measured by ELISA at recipient sites. The results are normalized to total insulin content and expressed relative to basal conditions (3, 2.8 and 3 mM glucose, respectively). (F) Non-ELISA based GSIS assay performed at Oxford University with 3 mM, 17 mM or 17 mM glucose with 20 nM exendin4 (ex4); n = 3-4 replicates of 10 clusters from 3 independent differentiations. (G) Non-ELISA-based GSIS assay performed at Imperial College, London, using both SC-islets and primary islets in the presence of 2.8 mM or 16.7 Glucose or 20 mM potassium chloride (KCl); SC-islets from n=4 bioreactors and primary islets derived from 5 donors. (H) ELISA-based GSIS assay performed at King’s College, London, with 3 mM versus 20 mM glucose dots represent technical replicates from n=2 bioreactors. (I, J) Cytosolic Ca^2+^ levels of (I) H1 (Fluo-8) and (J) H3 (Fura2) hESC-derived single SC-β cells in 3 mM glucose, 17 mM glucose and 10 mM KCl. SC-β cell selection was based on an increase of at least 20% in calcium levels from low to high glucose (n [H1] = 6-8 clusters from 3 independent differentiations; n [H3] = 6-11 clusters from 3 independent differentiations); Graphs show connecting lines and mean ± SEM. (K, L) β cell-specific Perceval-HR measurements showing ATP/ADP dynamics in response to 3 and 17 mM glucose at weeks 4 and 6 of Stage 7 differentiation. (K) Representative traces of SC-β cells transduced with the Perceval HR; Graphs show connecting lines and mean ± SEM. (L) Quantitative comparison of (left panel) the fold-change increase in response to glucose and (right panel) the area under curve (AUC) in 3 and 17 mM glucose (n [week 4] = 20 clusters from 4 independent differentiations, n [week 6] = 28 clusters from 6 independent differentiations). Statistical significance was tested using (C, E, H) a two-tailed unpaired Student’s *t* test with Welch’s correction, (D) Wilcoxon rank-sum tests, (F, L) a linear mixed-effects model with Tukey-adjusted post hoc tests to account for nested data structure, and (G) two-tailed paired Student’s *t* tests.

Sufficient functional maturity is essential for SC-islets to serve as an *in vitro* model for β-cell, obesity and diabetes research. Thus, we next assessed functional maturity by measuring glucose-stimulated insulin secretion (GSIS) performed at the production site (secretion index: 5.7 with IBMX, Figure 2E ‘Birmingham’). The cells were further benchmarked by alternative static GSIS protocols at three independent recipient sites post-shipment (Figures 2F-2H; ‘Oxford’, ‘Imperial’ and ‘King’s’). Static GSIS was used rather than dynamic GSIS, since this is a general technique available in most receiving research teams, and serves as an excellent control for function across sites. The cells consistently displayed glucose responsiveness across all recipient centres, despite differences in culture conditions, assays and experimenters.

Consistent with intact oxidative phosphorylation, SC-β cells within SC-islets demonstrated glucose-stimulated increases in ATP/ADP ratios, with enhanced metabolic activity observed in clusters at Stage7 week6 relative to week4, indicating further maturation in prolonged culture (Figures 2K–2L). Live-cell calcium imaging revealed glucose-induced rises in intracellular Ca²⁺ across the β-cell population in each batch examined, consistent with K_ATP_ channel closure, depolarisation and calcium influx through voltage-dependent Ca^2+^ channels (Figures 2I-2J). Together, these data demonstrate that SC-islets generated using our platform are glucose-responsive and exhibit key features of functional β-cells, including intact glucose-sensing, membrane depolarization and Ca^2+^ influx, and coordinated stimulus–secretion coupling, making them suitable for mechanistic studies of β-cell function and dysfunction in metabolic disease.

### Single-cell transcriptomics reveals β-cell–enriched populations with limited off-target heterogeneity dominated by EC-like cells

To define the cellular and transcriptional identity of cells generated using our platform, we performed single-cell RNA sequencing (scRNA-seq) on differentiated H1 and H3 derived SC-islets at Stage7 week2 (Stage7w2), which corresponds to maturing β-cells, and our timepoint of shipment (n=2 for each line). For the H1-derived cells, we specifically sequenced a high-performing (∼75% C-peptide+ cells by flow cytometry, replicate1), and a low performing (∼55% C-peptide+ cells, replicate2) production. In the high performing replicate (1), Uniform Manifold Approximation and Projection (UMAP) analysis reveals a relatively simple and well-resolved cellular landscape, consisting predominantly of three endocrine cell types: β-cells (69%), enterochromaffin (EC) like cells (24%), and α-cells (5%), annotated by marker gene expression (Figures 3A-3C). In the low-performing replicate (2), similar clustering was observed but with less favourable β-cell:EC-like cell ratios (Figure S3A). Thus, consistent with our flow cytometry–based estimates of cell identity (Figure S3B), scRNA-seq analysis indicates that β-cells constitute approximately ∼60-70% of the total cell population in an average SC-islet preparation, representing a high proportion of unsorted β-cells, making them ideal for research focused on β-cell biology, as well as genomics assays that primarily reflect the dominant cell-type. scRNA-seq analysis of SC-islets derived from a second stem cell line (hESC3, H3) revealed a slightly more heterogeneous cellular landscape with 57% β-cell, 24% EC-like cell, 15% α-cells, and a small population of somatostatin (SST) positive δ-cells (∼3%) (Figures S3C-S3F), with β-cell proportions in line with flow-cytometry based estimates (Figure S3D).

**Figure 3.**
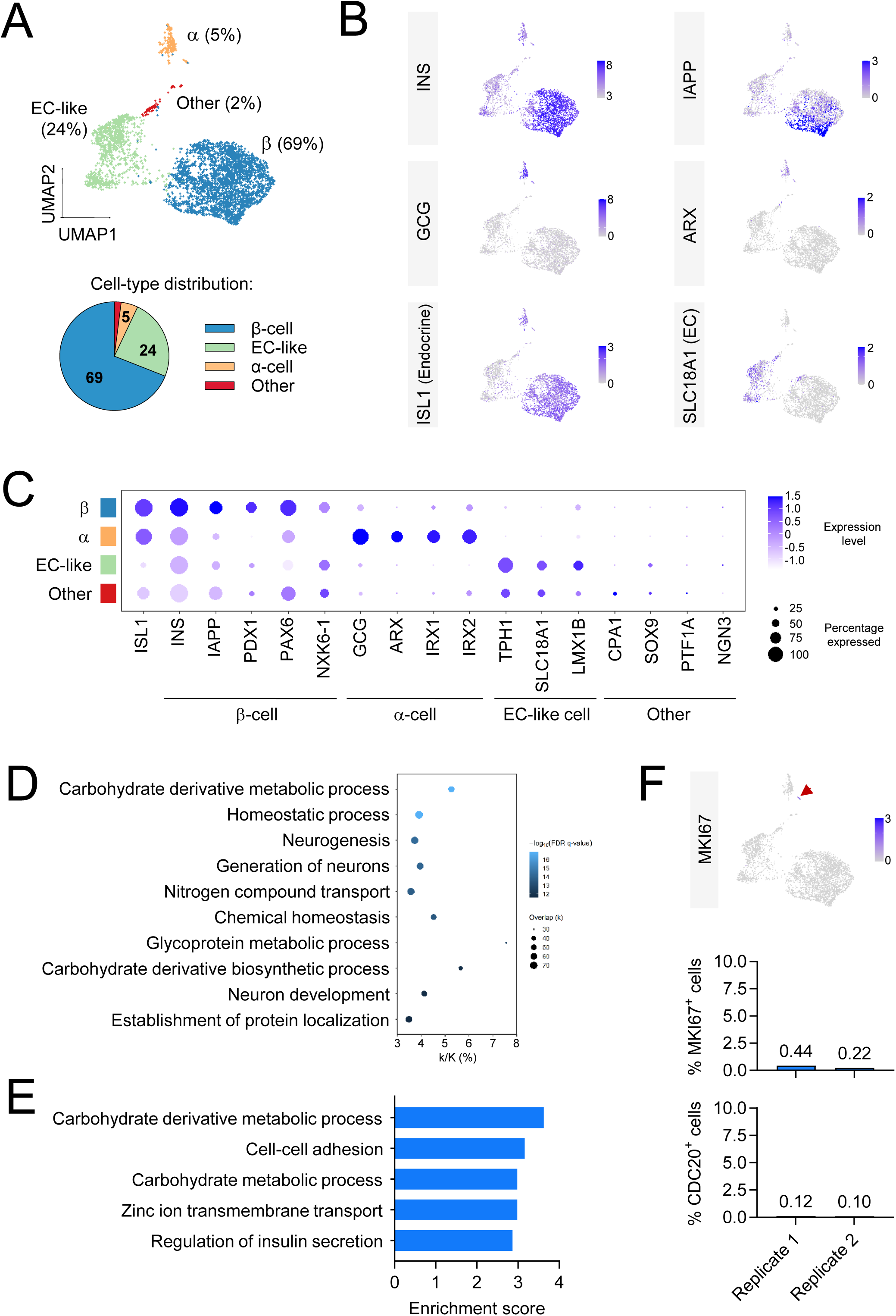
Transcriptomic landscape of SC-islets at single-cell resolution. (A) UMAP visualization of scRNA-seq data from SC-islets derived from the H1 hESC line at Stage7w2 (Replicate1). Lower panel: percentage of cells in each cluster. Clusters were annotated by cell type based on expression of islet hormone genes and established enterochromaffin (EC) cell markers. (B) Feature plots showing expression of key endocrine markers, including INS, IAPP, GCG, the α-cell transcription factor ARX, the pan-endocrine marker ISL1, and the EC-cell marker SLC18A1. (C) Dot plot summarizing average expression and the proportion of cells expressing marker genes associated with expected endocrine and off-target lineages across clusters. (D-E) Gene ontology analysis of top 500 marker genes enriched in Cluster 0 (β-cell cluster), performed by gene set enrichment analysis (GSEA, D) and DAVID bioinformatics tool (E), highlighting pathways associated with β-cell identity and function. (F) (Top) Feature plot showing expression of the proliferation marker MKI67 (red arrow). (Bottom) quantification of the proportion of cells expressing MKI67 or the alternative proliferation marker CDC20 across two independent Stage7w2 H1-derived SC-islet samples.

Gene ontology analysis of the top 500 marker genes (Table S3) for the β-cell cluster was enriched for terms associated with canonical β-cell function, including carbohydrate derivative metabolic processes, cellular homeostasis, zinc ion transport as well as insulin secretion (Figure 3D, E using DAVID^40^ and GSEA^41^ respectively). In line with previous studies, EC-like cells displayed a more prominent neuronal transcriptional programme, while the small α-cell population displayed gene ontology terms consistent with hormone production (Figure S3G).

A key limitation of this *in vitro* model is the presence of off-target lineages that might complicate experimental interpretation. As previously reported, the main contaminant cell type in both H1 and H3 derived SC-islets comprises EC-like (enterochromaffin-like) cells, a serotonin-producing endocrine lineage marked by SLC18A1 (Figures 3B-3C). In H1-derived SC-islets, markers for pancreatic progenitor (NGN3), ductal (SOX9, HNF1B, MUC1), exocrine (CPA1, PRSS1), mesenchymal (VIM, COL1A1, TAGLN), and endothelial (PECAM1, VWF, EMCN) populations are either expressed at <10% or at levels similar to primary-β-cells (Table S4, Figures 3C and S3H), indicating a high degree of endocrine purity. Consistent with this, newly identified^42^ non-endocrine *in vitro* markers associated with adverse *in vivo* outcomes, ANXA2, FN1 and YAP were detected in only 1-2% of cells and were not restricted to a particular cell type, neither did they co-localize in the same cells (Table S4 and data not shown). Proliferative capacity was low; with only a small fraction of cells (<0.5%) expressing cell-cycle markers (MKI67, CDC20; Figures 3F and S3I) and co-expressing α-cell markers ARX and GCG, consistent with a minute proliferative α-cell population also present in donor derived human islets^43^ (Figure S3I).

To further assess heterogeneity and alternative lineage presence within the β-cell population, we examined a broader list of alternative lineage markers at multiple expression cut-off values in our H1 and H3-derived SC-β cells and compared them to three primary β-cell samples^44–46^ using scRNA-seq (Table S4). Our results show that the majority of alternative lineage markers in SC-β cells fall within the expression range observed in donor-derived primary β-cells, with the exceptions of KRT19 (ductal), FHOD3 (cardiac) and EC-like/neuronal markers. In the absence of additional cardiac or ductal markers, the upregulation of KRT19 and FHOD3, both associated with intermediate filament organization, may reflect spinner culture-related effects, where cell cohesion and cytoskeletal reinforcement are essential^47^.

Collectively, these findings establish a robust, endocrine-focused *in vitro* platform with defined and minimal contaminant populations, enabling the study of β-cell and islet function without confounding from off-target cell types.

### SC β-, α-, and δ-cells closely resemble primary islet cell types but retain distinct transcriptional features

To evaluate the transcriptional fidelity of SC-islet cell types relative to their primary counterparts, we analyzed scRNA-seq data from our (H1 and H3) SC-islets and three donor-derived human islet samples^45–47^. We performed guided clustering of all samples and subclustered α, β, and δ cell types based on marker expression (Figures 4A-4C). We then assessed transcriptional similarity by specific interrogation of the top most different genes (highly variable features, VF) across all clusters as well as the specific VFs belonging to endocrine cells. This analysis, with different sets of VFs and with aggregated or averaged expression (Figures 4D and S4A-E), reveals a striking transcriptional similarity between primary/SC-β cells (Pearson’s *r*^2^ = 0.9-0.98), primary/SC-α cells (*r^2^*=0.64) and primary/SC-δ-cells (*r^2^*=0.97). In contrast, all three islet cell types (SC and primary) appear transcriptionally distinct from a primary non-endocrine cell type (ductal) (Figures 4D and S4A-E). We obtain similar results using an alternative clustering algorithm (Figure S4F-I, Methods). This transcriptional similarity, coupled to high endocrine purity, enables more accurate genomic studies focusing on the human β-cells.

**Figure 4.**
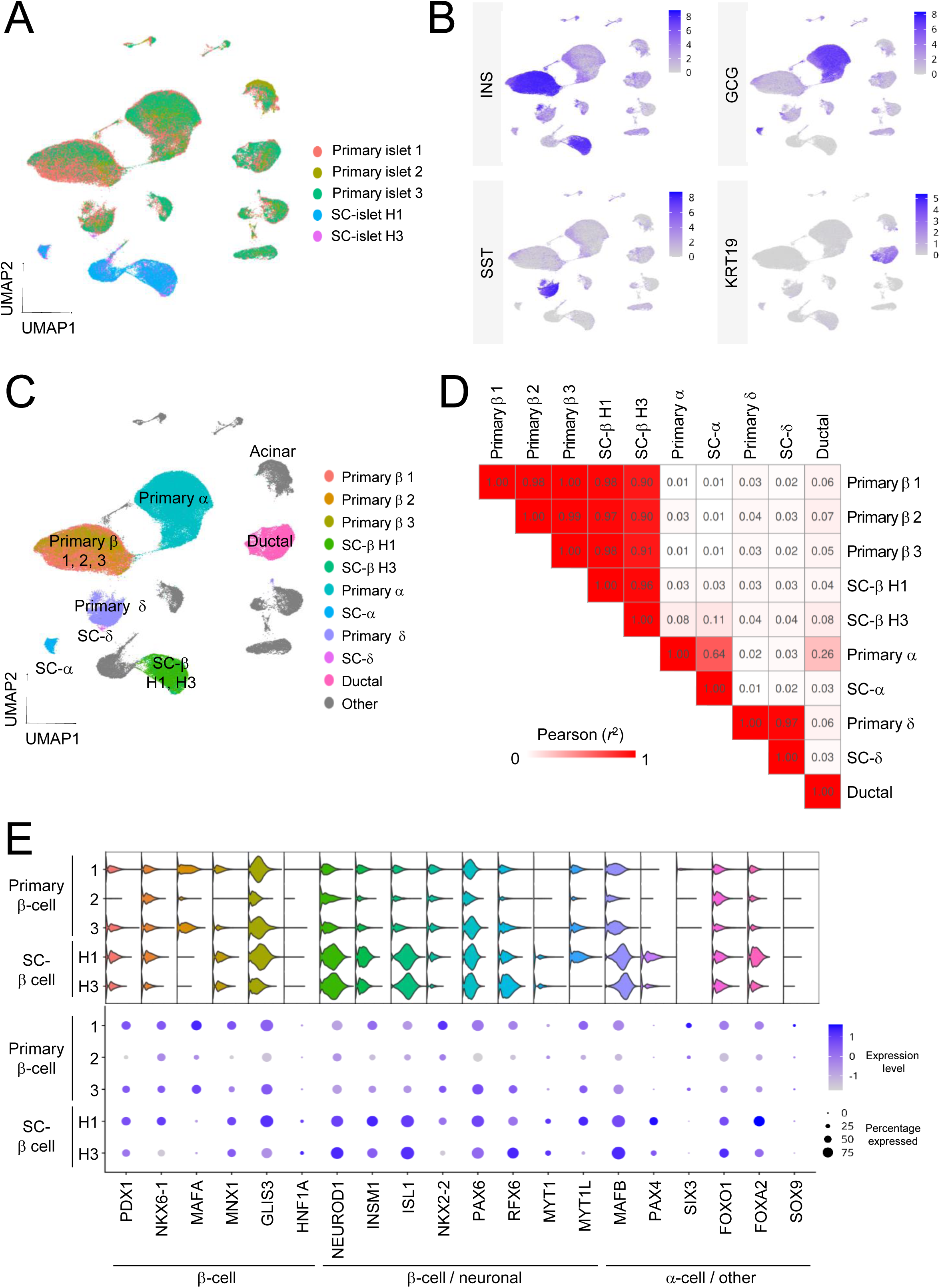
Transcriptional cross-comparison between primary and SC-islet cell types. (A) Harmony integrated UMAP visualization of scRNA-seq from three publicly available healthy primary-islet datasets, and SC-islets derived from H1 and H3 hESC lines. Clusters were annotated by dataset. (B) Feature plots showing expression of key endocrine markers, including INS, GCG, SST, and ductal cell marker KRT19 from (A). (C) Cell types annotated based on dataset and marker expression (D) Heat map of the Pearson correlation (*r*²) matrix calculated by comparing the aggregated RNA counts of each cell population‘s expression of the top 3000 HVFs computed from all cells in the entire dataset (all correlations BH adjusted p-value < 0.001). (E) Panel of violin plots (top) and a dot plot (bottom), grouped by β-cell sub-populations, showing log-normalized RNA counts of key marker genes.

Previous studies described an abnormal neuroendocrine transcriptional programme expressed in SC-β cells in comparison to primary human β-cells and islets^4,22,23,48^. Targeted interrogation of scRNA-seq data from primary and SC-β cells reveals a similar upregulation of neuronal gene pathways as well as key neuronal transcription factors shared between the endocrine pancreas and neurons, such as ISL1, NEUROD1 and PAX4 in our SC-β cells (Figure 4E, GSEA Analysis, Table S5). Interestingly, an abnormal upregulation of a neuronal programme is not necessarily associated with downregulation of genes related to β-cell specific functions; genes previously identified as associated with clustered islet enhancers^21,22^ known to drive cell identity programmes or hallmark β-cell genes were not depleted in SC-β cells as assessed by GSEA (Table S5). These findings indicate successful establishment of β-cell identity in SC-β cells but point to an inability to efficiently repress alternative lineage transcriptional programmes that may interfere with function.

Collectively, our results suggest that SC-islet cell types are transcriptionally distinct from human donor derived islets but maintain gene regulatory pathways that are sufficiently similar to allow modelling of β-cell function and disease-relevant mechanisms.

### Functional metabolic deficits in SC-islets occur in the absence of major transcriptional alterations

Previous studies reveal that stem cell derived β-cells are metabolically immature^6,23^. Our SC-islets also remain inferior to human donor derived islets in their glucose stimulated insulin secretion responsiveness (Figures 2 and S2). We examined whether the functional or metabolic immaturity is a result of transcriptional defects in SC-islets due to incomplete expression of genes related to metabolic pathways. To do this, we compared gene expression profiles of our SC-islets (H1 and H3, n=6) with those of high-purity (n=12, INS:CPA >5 and INS/KRT19 ratio > 15) human donor derived islet samples using bulk RNA-sequencing, which provides greater read depth for more robust pathway analysis. Differential gene expression analysis followed by gene ontology analysis of dysregulated genes suggests upregulation of neuronal gene pathways in SC-islets as previously reported^4,22,23,48^ (Figures S5A-S5B). We also observe downregulation of vasculature and tube development pathways in SC-islets (Figures S5A and S5C), in line with the fact that *in vitro* differentiated organoids are devoid of pancreatic ducts and vasculature.

Interestingly, only minor differences are observable in the expression levels of genes involved in the TCA cycle, malate, aspartate, pyruvate and glutathione metabolism, with none of the metabolic pathways significantly downregulated in SC-islets in comparison to donor derived islets as assessed by GSEA analysis (Figure S5D; Table S5). Moreover, genes belonging to insulin secretion or glycosylation pathways are not significantly dysregulated in SC-islets when compared to primary islets (Table S5). We observe similar outcomes for transcriptional regulation of these pathways in SC- and primary β-cells based on scRNA-seq data with only mitochondrial genes modestly downregulated in SC-β cells, compared to primary β-cells (Table S5). These data show that, despite intact metabolic pathway gene expression at the transcriptomic level, reduced metabolic flux and inefficient coupling to ATP production constitute the principal bottleneck in glucose-stimulated insulin secretion in SC-islets.

## MATERIALS AND METHODS

### Cell lines and sources

Human embryonic stem cell (hESC) line H1 was sourced from WiCell (USA), hESC3 (H3) from Ed Stanley (Murdoch Children’s Research Institute, Australia). Ethical approvals were obtained from the UK Medicines and Healthcare Products Regulatory Agency (MHRA) for research using these lines (SCSC24-33, SCSC24-34).

### Human Islets

Isolated human islets from cadaveric donors were provided by the Integrated Islet Distribution Program (IIDP) and cultured in RPMI with 5.5 mM glucose, 10% FBS, 2 mM L-glutamine, 37°C/5% CO_2_. This study used data from the Organ Procurement and Transplantation Network (OPTN). The OPTN data system includes data on all donor, wait-listed candidates, and transplant recipients in the US, submitted by the members of the Organ Procurement and Transplantation Network (OPTN). The Health Resources and Services Administration (HRSA), U.S. Department of Health and Human Services provides oversight to the activities of the OPTN contractor. Internal ethical approvals were obtained to work with human tissue at Imperial College.

### Stem cell maintenance and differentiation into stem cell derived β-cells

Human pluripotent stem cells (H1 and H3) were maintained on Geltrex (Thermo Fisher, 12053569) in mTeSR Plus medium (STEMCELL Technologies, 100-1130) and StemFlex (Fisher Scientific, 15627578) respectively and passaged using ReLeSR (STEMCELL Technologies, 100-0484) at ratios of 1:10 (H1) and 1:18 (H3) following manufacturer’s instructions. Cells destined for differentiation were maintained in culture for a minimum of 6 weeks with strict 24-hour interval medium changes, followed by cryopreservation (10% DMSO, Sigma-Aldrich D2438; 90% EmbryoMax ES Cell Qualified FBS, Sigma-Aldrich ES-009). Cryopreserved cells were thawed and maintained in culture for 2 weeks prior to differentiation. Differentiation to stem cell-derived β-cells (SC-islets) was performed as described^7^ with the following modifications; 13 × 10⁶ H1 or 10 × 10⁶ H3 cells were seeded per 10 cm dish 24 h prior to induction. Definitive endoderm was induced using 110 ng/ml Activin A (Qkine, Qk001-1000) and 3 μM CHIR99021 (STEMCELL Technologies, #72054) in 15 mL medium per dish; subsequent stages used 10 mL per dish. Stage 2 included KGF (Qkine, QK046; 50 ng/mL) for 3 days, and Stage 4 included EGF (Qkine, Qk011-1000; 100 ng/mL). Cells were transferred to AggreWell plates (STEMCELL Technologies, 34425) on day 10 and, after 24 h, moved into ABLE 30 mL bioreactors spinning at 60 rpm (ABBWBP03N0S-6) at 30 mL per 10 cm dish equivalent. Stage 7 medium contained zinc sulfate (10 μM), heparin (10 μg/mL), and ZM447439 (S1103; 0.5 μM), with additional daily pulses of 1 μM ZM447439 for 3 consecutive days during week 2. phASC was included at 50 μg/mL during Stage 7. Stage 7 media was changed every other day, without exception. When needed, cells were transported to other centres in full 15 mL tubes with less than 4 h transit time using warm-packs at 37° C. At recipient sites, the cells were maintained in 6-well dishes, shaking at 100 rpm.

### Flow cytometry analysis

Cells were dissociated into single-cell suspensions using TrypLE Express (Gibco, 12605-028) or Accutase cell dissociation reagent (Gibco, A1110501) for 5–8 min at 37°C, followed by fixation in IC Fixation Buffer (Invitrogen, 00-8222-49) for 20 min. Cells were washed with PBS, blocked in blocking buffer CAS block (Invitrogen, 008120) with 0.2% Triton-X100 (Sigma,1003569923), for 1 h on ice, and incubated overnight with primary antibodies: anti-C-peptide (clone U8-424, 1:150; BD Biosciences, 565831), anti-glucagon (U16-850, 1:100; BD Biosciences, 565891), anti-ISL1 (Q11-465, 1:100; BD Biosciences, 562547), anti-NKX6-1 (R11-560, 1:100; BD Biosciences, 567608), anti-PDX1 (658A5, 1:100; BD Biosciences, 562161), anti-SOX17 (P7969, 1:200; BD Biosciences, 562205), and anti-FOXA2 (N17-280, 1:200; BD Biosciences, 561589). Cells were washed once with PBS, resuspended in PBS and run on a CytoFLEX flow cytometer (Beckman Coulter). Data were acquired and analyzed using CytExpert 2.7 software.

### Glucose stimulated insulin secretion assays at production and recipient sites

GSIS was performed as described^49^ with the following modifications: approximately 50-100 clusters of SC-islets were transferred to starvation medium consisting of Stage7 media components in no glucose RPMI medium (Fisher Scientific, 11470425) and incubated at 37°C, 5% CO_2_ for 1 hour, shaking at 60 rpm. Static GSIS was measured by incubation in Krebs-Ringer buffer^49^ containing 2 mM or 16 mM glucose and 0.5mM IBMX (Merck, I5879) for 30 minutes at 37°C. SC-islets were lysed in TETG buffer containing 20 mM Tris-HCl (pH 8.0), 1% Triton X-100, 10% glycerol, 137 mM NaCl, and 2 mM EGTA, prepared in distilled water, and supplemented with EDTA-free protease inhibitor cocktail (Merck, 1183617001). Cell lysate was sonicated using a Bioruptor sonicator (Diagenode) set to medium power, with 30s ON, 30s OFF for 5-6 cycles at 4°C. Total protein content was measured using the Pierce BCA protein assay kit (ThermoFisher Scientific, A55864). Insulin content of supernatant and cell lysate was measured using a Human Insulin ELISA (Merck, EZHI-14K).

GSIS at recipient sites were performed at (i) Imperial College, London, as described for human islets and SC-islets^7,50,51^ using ultra-sensitive insulin homogeneous time-resolved fluorescence (HTRF) assay kits (Cisbio) and a PHERAstar Plus microplate reader (BMG Labtech); (ii) Oxford University performed as described^52^ with exendin4 at 20 nM; (iii) King’s College, London, as described^53^.

### Ca^2+^ imaging or ADP/ATP ratio

Ca^2+^ and ATP/ADP live cell imaging of SC-islet clusters was performed at 34 °C using a CrestOptics X-Light V2 spinning disk coupled to a Nikon Ti-E microscope, 89 North LDI-7 Laser Diode Illuminator, Prime BSI Express sCMOS and 10x/0.8 NA objective.

The Ca^2+^ dye Fluo-8 (AAT Bioquest, 21082) was complexed with pluronic acid (20% in DMSO) at 42 °C for 5 min. 10-20 SC-islet clusters were incubated with 10 µM of Fluo-8 for 1 h in RPMI 1640 supplemented with 10% FBS and 1% P/S at 37°C and 5% CO2. SC-islet clusters were then washed twice and incubated for 15 min in HEPES-bicarbonate buffer containing in mM: 120 NaCl, 4.8 KCl, 24 NaHCO_3_, 0.5 Na_2_HPO_4_, 5 HEPES, 2.5 CaCl_2_, 1.2 MgCl_2_, and 3mM D-glucose. SC-islets were imaged with an excitation wavelength of λ = 470 nm and emission was recorded at λ = 500-550 nm every 5 s. After 10 min of recording, the glucose concentration was increased to 17 mM. At the end of the experiment, 10 mM of KCl was added as a positive control. In experiments using H3 SC-islets, Fura2 (Hello bio., HB0780) was used in place of Fluo8 because of spectral overlap with GFP. Dye loading was performed as described for Fluo8, with excitation at λ = 340 and 385 nm delivered using a FuraLED system and emission detected at λ = 470–550 nm. Single-cell Ca^2+^ analysis was performed using a custom-written ImageJ macro to ensure an unbiased approach. For Fluo-8, the Ca^2+^ traces were normalized as F/F0 where F = fluorescence at a given timepoint, and F_0_ = fluorescence at baseline. For Fura2, Ca^2+^ traces are presented as the ratio of 340/385 nm, then normalized as above.

For ATP/ADP measurements, SC-islet clusters were transduced for 48-72 h with the beta cell-specific FRET sensor Ad-RIP-Perceval-HR^54,55^ (a kind gift from Prof. Matthew J. Merrins, Yale University), prior to imaging as above. Perceval-HR was excited sequentially at 405 nm and 470 nm, and emission collected at λ = 500–550 nm every 3 seconds. After 2 minutes of recording, the glucose concentration was increased to 17 mM, as indicated. ATP/ADP ratios were calculated as the 470/405 nm excitation ratio, before normalising as R/R_0_ where R = 470/405 ratio at a given timepoint, and R_0_ = 470/405 ratio at baseline.

### Bulk and single-cell RNA sequencing

Approximately 1 × 10⁶ SC-islets were harvested in 1 mL TRIzol Reagent (Invitrogen, 15596026), and total RNA was extracted followed by DNase treatment (Sigma, D5307-1KU) according to the manufacturer’s instructions. Bulk RNA sequencing was performed by Novogene (Cambridge, UK) using an Illumina platform (NovaSeq6000), generating 150 bp paired-end reads at a depth of ∼30 million reads per sample.

For single-cell RNA sequencing, SC-islet clusters were dissociated using 1:1 mixture of TrypLE Select (Gibco, 12563-029)) and Trypsin-EDTA (Sigma, T4174; 10X stock diluted 1:10 with PBS) for 10 min at 37°C. The reaction was quenched with5% FBS-PBS, and cells were washed twice in cold 0.04% BSA-PBS solution. Dissociated cells were passed through a 30 μm strainer prior to processing using the 10x Genomics Chromium platform at the University of Birmingham Genomics Facility.

### Transcriptomic data analysis

Single-cell RNA-seq data were aligned to the human reference genome (hg38) using Cell Ranger (10x Genomics), and downstream quality control and analysis were performed using Seurat with default parameters. Cells failing quality control or exhibiting negligible expression of MALAT1 (a nuclear-retained lncRNA, indicative of empty droplets) were excluded. Feature plots, dot plots, and violin plots were generated using Seurat, while heatmaps and bubble plots were produced using ggplot2 in R (v4.4) ClusterProfiler, Gene Set Enrichment Analysis or DAVID were used for gene ontology analysis. Following quality control filtering, cell numbers were as follows: SC-islet H1 replicate 1 (n = 4,406) and replicate 2 (n = 4,586), and SC-islet H3 replicate 1 (n = 3,005) and replicate 2 (n = 3,554). For downstream analyses, H1 replicate 1 and both H3 replicates were included. H1 replicate 2, low-performing differentiation, was analyzed separately for validation purposes.

Published bulk RNA-seq data were analyzed as previously described^21^. 12/208 human islet samples met purity thresholds (INS/CPA > 5 and INS/KRT19 > 15) and were used for this article. Data were obtained from GEO/SRA (SRR12885670, SRR6376127, SRR12885680, SRR12885672, SRR6376140, SRR12885691, SRR6376164, SRR6376182) and the EGA (EGAD00001003992).

For Figure 4 and S4, three primary-islet scRNA-seq datasets were selected that had healthy, untreated samples and were all sequenced with 10X Genomics. Healthy samples for each dataset^44–46^ were downloaded from GEO with accession numbers GSE221156, GSE148073, and GSE217837, and aligned the same way as SC-islet samples.

R (v4.3.3) on RStudio (v2026.01.0+392) was used to process count matrices using Seurat^56^ (v5.4.0). Count matrices were grouped by sequencing libraries and then filtered for strict quality control (QC) in 3 steps. First, SoupX^57^ (v1.6.2) was used to automatically estimate ambient RNA and adjust counts to reduce contamination. If contamination could not be estimated, then the contamination fraction was set to 20%. Second, only cells with at least 200 features and features in at least 3 cells were kept for further analysis. Cells within ±2.5 Mean Absolute Deviations (MADs) of RNA nFeatures and nCounts with lower thresholds clamped at 0, and with cells <20% mitochondrial reads were selected. Lastly, DoubletFinder^58^ (v2.0.6) was used to estimate doublets using default settings. The doublet rate was set to 0.75% per 1000 cells recovered, in accordance with the 10X user guidelines, and classified doublets were removed. QCed sequencing libraries were then unified for each dataset and any remaining cells exhibiting low MALAT1 expression were removed^59^. The same QC steps were taken for SC-islet datasets. Datasets (primary-islet 1, 2, 3 and SC-islet H1, H3) were log-normalised separately and then scaled together. Linear dimension reduction was performed, and the standard deviations of the principal components were plotted to determine a suitable number of components to compute nearest neighbours. Unsupervised clustering was performed and Uniform Manifold Approximation and Projection (UMAP) coordinates were calculated. Resulting clusters were visualised and any presenting with characteristics of empty droplets (i.e. multi-hormonal expression and consistently low nFeatures) were removed, then remaining cells were renormalised per dataset and scaled together. After this QC, cell numbers were as follows: primary-islet 1 (n = 79,771), primary-islet 2 (n = 12,063), primary-islet 3 (n = 18,702), SC-islet H1 replicate 1 (n = 3,748) and replicate 2 (n = 3,741), and SC-islet H3 replicate 1 (n = 4,146) and replicate 2 (n = 3,659).

Principle components were recalculated so datasets could be integrated using Harmony^60^, then nearest neighbours, unsupervised clustering, and UMAP coordinates were recomputed. As an alternative method to integration, after removing clusters composed of empty clusters, datasets were combined and SCTransformed together (Figure S4F-I). This was tested as all datasets were composed of similar cell types and sequenced with the same platforms (10X v2/3). Datasets appeared to cluster appropriately, with cells from primary-islets clustering together (figure S4F) and with their correct cell types.

Pancreatic and endocrine markers, beta (*INS*, *IAPP*), alpha (*GCG*, *ARX*), delta (*SST*), epsilon (*GHRL*), gamma (*PPY*), enterochromaffin (*TPH1*), ductal (*KRT19*), acinar (*PRSS1*), vascular (*PECAM1*), immune (*PTPRC*), and stromal (*COL1A1*) were visualised, and cell types were labelled using their respective markers.

Differential gene expression analysis between SC- and primary β-cells was performed using default settings on Seurat.

Pearson correlation coefficients of determination (*r*²) were calculated to compare primary-islet cell types and their stem-cell derived counter parts. Ductal and acinar cells were included as non-endocrine controls. Highly Variable Features (HVFs) were selected for various cell populations by taking genes with the largest residual variance from a regularized negative binomial regression model calculated per dataset, then common HVFs across datasets were extracted. HVFs were computed for the following cell populations: all cells, primary and SC-islet endocrine cells (α, β, γ, δ and ε), primary and stem-cell derived β cells (data not shown), all cell types separately then combined (data not shown), and all cell types down sampled to equal sizes (data not shown).

To test transcriptional similarity between primary and stem-cell-derived beta cells, RNA expression values for cell populations were summarized by two methods. The first method aggregated the expression of features per cell population by summing their raw counts, and the second method averaged the expression of features per cell population by exponentiating log-normalised values then calculating the mean in non-logged space. Pearson correlation coefficients (r) and Benjamini-Hochberg (BH) adjusted p-values were calculated by comparing the summarized expression profiles of HVFs of varying cell populations and coefficients of determination (r²) were visualised. Pearson correlations (r) and BH adjusted p-values were computed using the psych R package (v2.6.3), and correlation matrices were visualised using the pheatmap R package (v1.0.13).

### Statistical and variability analysis

All statistical analyses were performed with GraphPad Prism 11 (version 11.0.0) or R (v4.5.3) using the indicated methods. Normal distribution was tested using Anderson–Darling, D’Agostino & Pearson and Shapiro–Wilk tests. A p value less than 0.05 was considered significant: ns, not significant; *p < 0.05; **p < 0.01; ***p < 0.001; ****p < 0.0001. Unless otherwise stated, all bars represent mean ± SEM.

Variability between batches and within individual batches was assessed using the coefficient of variation (CV), calculated as the standard deviation divided by the mean, expressed as a percentage.

### Data access

RNA sequencing data has been deposited at GEO Data (GSE330107), please find the reviewer’s access below. To review GEO accession GSE330107: Go to https://www.ncbi.nlm.nih.gov/geo/query/acc.cgi?acc=GSE330107 Enter token anajyuqanduhfmf into the box

## DISCUSSION

Despite substantial investment, sizable challenges remain for T1D regenerative therapies^24,25^ and addressing these challenges will require collaborative efforts from researchers specializing in various fields such as vasculature, bioengineering, materials science, and immunology. Access to high-quality SC-islets remains a critical bottleneck, not only for the advancement of regenerative therapies but also for the broader diabetes and pancreatic biology research community. Here, we describe the standardization of an SC-islet production protocol in a facility setting and the transcriptomic and functional characterization of the resulting cell clusters across multiple recipient sites. Our platform combines controlled stem cell culture, improved endoderm induction, aurora kinase inhibition to limit proliferative cells, and scalable bioreactors to enhance culture uniformity. These refinements increase consistency and β-cell yield (∼70% C-peptide^+^/ISL1⁺), while minimizing off-target and bihormonal cells. Recently published protocols also achieve similar β-cell proportions using modified protocols^18,26,27,61,62^, with some deploying mid-scale bioreactors^18,26,29^. Taken together, our platform provides a robust and accessible SC-islet model *on par* with the most recent protocols, available through an academic, non-profit facility^63^.

For SC-islets produced using our platform, a high proportion β-cell content and a profoundly similar transcriptional programme to their primary equivalents does not translate to an equivalent function to the human islet (Figures 2 and S2). We hypothesize that various factors may contribute to this. First, we observe a lower SC-α cell content in SC-islets compared to primary islets (5-15% vs 30-40% respectively). For a human islet, correct cellular composition and spatial organization is needed for intra-islet signalling^64^ and *in vivo* studies support the notion that enabling paracrine signalling improves SC-β cell function^42,65^. The data is less clear on *in vitro* function with one study reporting no impact of SC-β/α cell ratios on SC-islets function^41^. We note that this discrepancy may be due to the subpar functionality of SC-α cells *in vitro*, potentially maturing *in vivo*. In our hands, SC-α cells appear to show the greatest transcriptional divergence from their primary counterparts among the islet cell subtypes (Figure 4D) and that their suboptimal function *in vitro* might be negligible. Second, functional assessments for our SC-islets were primarily conducted at Stage 7 (week2-3, close to time of shipment), which is the optimal window for functional studies in the original protocol; however, our SC-islets continue to improve functionally later in Stage7 (weeks 4–6, Figure 2J). Finally, despite a high similarity between primary and SC-β cells’ transcriptional landscape, obvious differences in the expression of gene pathways and key genes, such as INS and MAFA^66^ remain. We note that despite low levels of MAFA mRNA in our SC-islets, MAFA promoter remains accessible, as assessed by Assay for Transposase-Accessible Chromatin using sequencing (ATAC-seq) of mature SC-islets (Figure S6A), suggesting that absence of MAFA mRNA might reflect a [missing] signal dependent post-transcriptional regulation rather than incomplete establishment of β-cell identity. This highlights the need for signalling inputs that maintain β-cell function after identity has been established. Our data, together with that of others, suggest that there remains scope to further refine differentiation protocols to achieve more mature β-cells, including by better repression of alternative lineage (neuronal and α) transcriptional programmes and further enhancing β-cell specific pathways associated with functional maturation *in vitro*.

In summary, despite their limitations, the functionality, high purity and transcriptional fidelity of SC-islets generated using our platform make them well suited for a broad range of downstream applications. The predominance of β-cells ensures that genomic and functional readouts are not confounded by heterogeneous cell populations or bihormonal cells, while the expression of key T1D antigenic peptides (Figure S6B) supports their use in immunological assays, an important area of focus in cell replacement therapies for T1D. Importantly, SC-islets retain functional signalling pathways, making them amenable to drug screening and studies relevant to diabetes and obesity. The SC-islets have already been integrated into organ-on-chip systems and used to generate targeted knockout models (e.g. CFTR, ALMS1, data not shown), further highlighting their versatility. Collectively, these features establish our platform as a robust and accessible *in vitro* system for non-specialist researchers seeking to interrogate human β-cell biology across diverse experimental settings.

## Acknowledgements

This research was supported by UKRI MRC Impact Acceleration Account block grant, grant (MR/X502996/1), NC3Rs grant (APP47224), the Birmingham Fellowship Programme, Diabetes UK RD Lawrence Fellowship (20/0006136), and an Academy of Medical Sciences Springboard Award (SBF006\1140) to I.A as well as the academic facility BetaCell Birmingham. D.J.H. was supported by Diabetes UK (22/0006389) and UKRI ERC Frontier Research Guarantee (EP/X026833/1) Grants and funded by the Bukhman Centre for Research Excellence in Type 1 Diabetes. This work was supported on behalf of the “Steve Morgan Foundation Type 1 Diabetes Grand Challenge” by Diabetes UK and SMF (grant number 23/0006627 to I.A. and D.J.H.). A.H.S. was supported by a Novo Nordisk – Oxford Fellowship. Human pancreatic islets were provided by the NIDDK-funded Integrated Islet Distribution Program (IIDP) (RRID:SCR_014387) at City of Hope, NIH Grant # 2UC4DK098085 and a JDRF-funded IIDP Islet Award Initiative to S.J.M. (BS643Pi) as well as a Diabetes Research & Wellness Foundation (DRWF) project grant awarded to S.J.M. The research was funded by the National Institute for Health Research (NIHR) Oxford Biomedical Research Centre (BRC). The views expressed are those of the author(s) and not necessarily those of the NHS, the NIHR or the Department of Health. The project involves an element of animal work not funded by the NIHR but by another funder, as well as an element focussed on patients and people appropriately funded by the NIHR. Some human islet data reported here have been supplied by UNOS as the contractor for the Organ Procurement and Transplantation Network (OPTN). The interpretation and reporting of these data are the responsibility of the author(s) and in no way should be seen as an official policy of or interpretation by the OPTN or the U.S. Government.

We sincerely thank Timo Otonkoski, Hazem Ibrahim, Caroline Gorvin, Holger Russ, David Adams, Wiebke Arlt, Martin Hewison, Dan Tennant, Tristan Rodriguez Shanta Persaud and Adriana Flores-Langarica for reagents and advice. We acknowledge the support of the Genomics Birmingham Technology Hub and the Flow Cytometry Platform at the College of Medicine and Health, University of Birmingham, for access to equipment and technical expertise.

---

**Figure S1.**
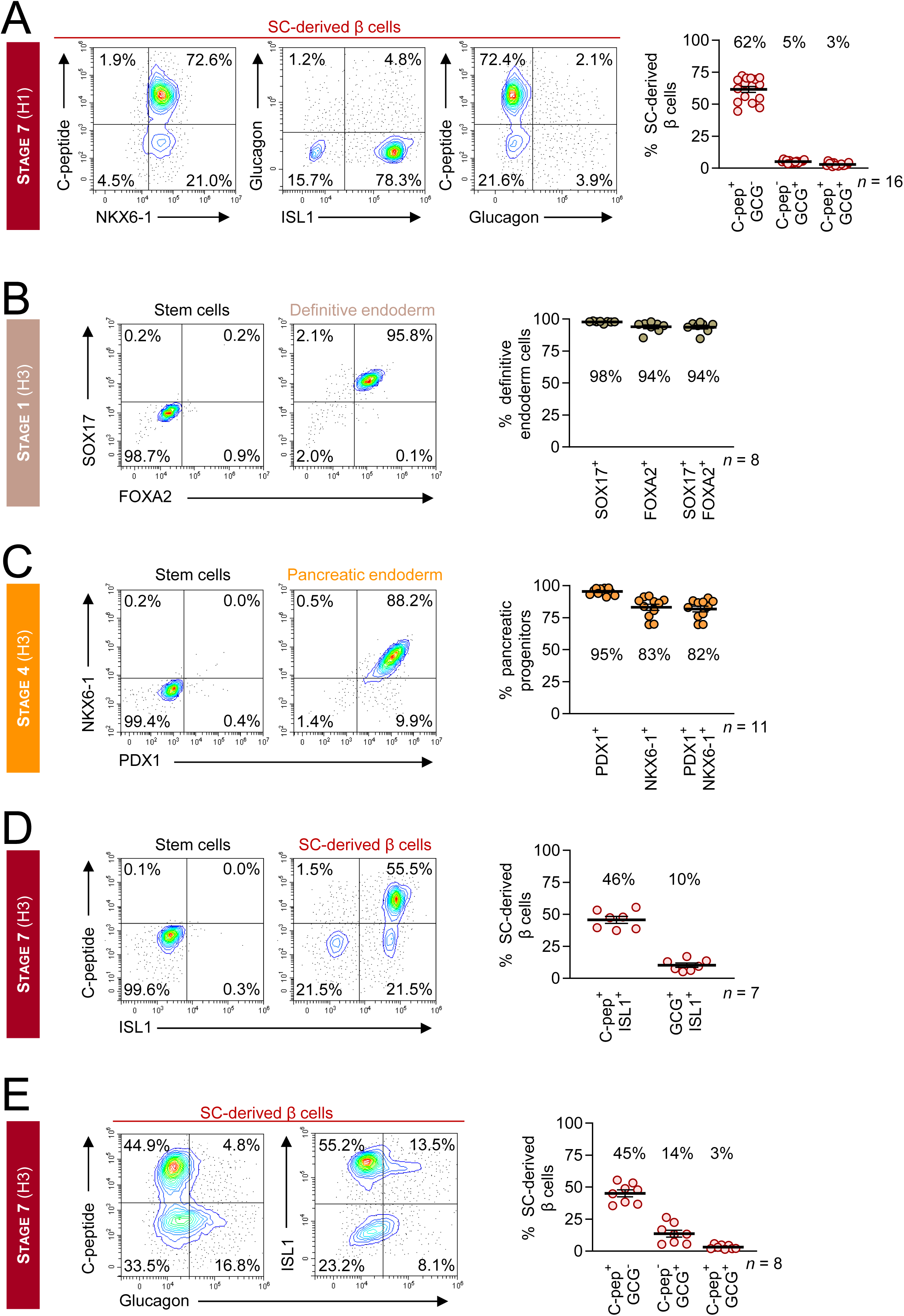
SC-islet identity marker analysis by flow cytometry. (A) Representative flow cytometry contour plots (left panels), and quantification across independent differentiations (right panel) for C-peptide/NKX6-1 or Glucagon/ISL1 or C-peptide/Glucagon (poly-hormonal) double positive cells in Stage7w2 SC-islets derived from H1 hESC line. (B-E) Representative flow cytometry contour plots (left panels), and quantification across independent differentiations (right panels), showing marker expression for Stage7w2SC-islets (B) definitive endoderm (Day3, FOXA2/SOX17), (C) pancreatic progenitor (Day12, PDX1/NKX6-1), and (D, E) mature endocrine stages (Stage7w2SC-islets, Day37, ISL1/C-peptide; NKX6-1/C-peptide; ISL1/Glucagon; C-peptide/Glucagon: poly-hormonal cells or ISL1/Glucagon: α-cells) derived from hESC3 (H3) line. Mean percentages of (co-)expression are shown by the bars, with the exact values annotated.

**Figure S2.**
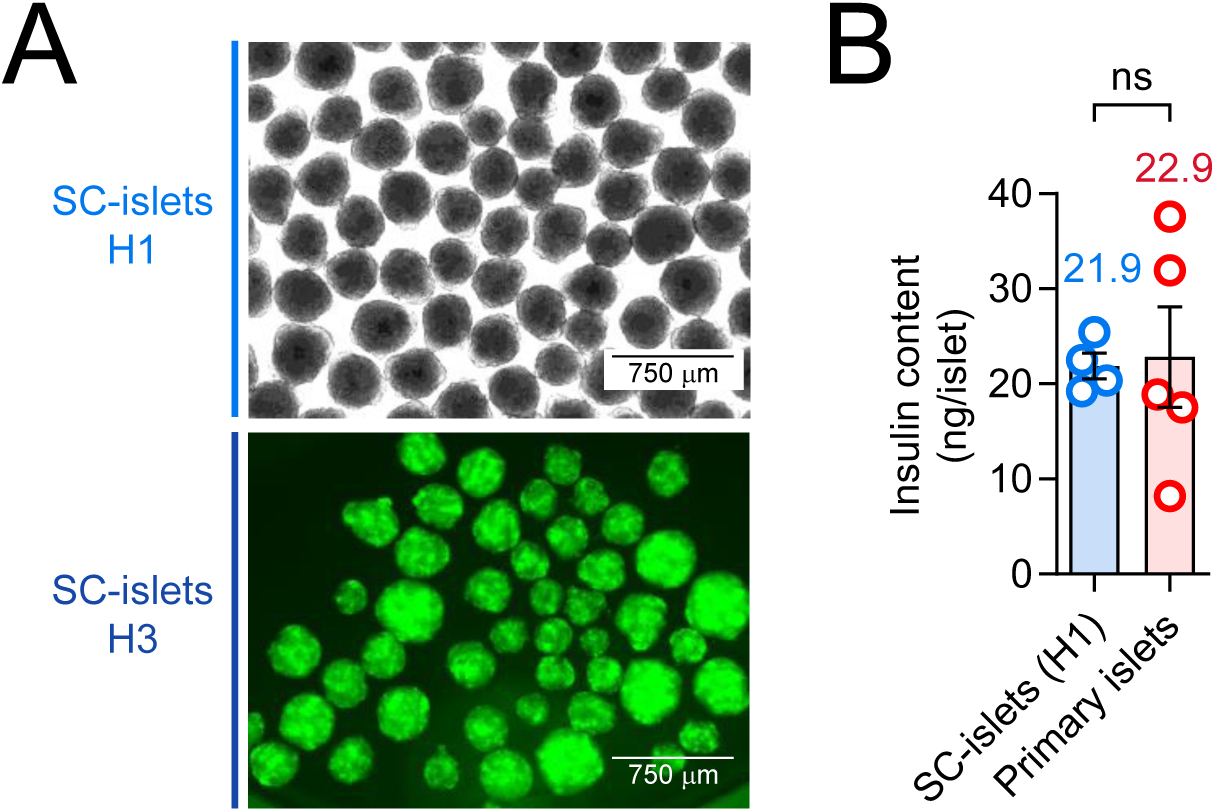
Comparable insulin expression in SC-derived and primary islets. (A) Microphotographs of (top panel) H1-derived and (bottom panel) H3-derived SC-islets. The green fluorescence reveals the expression of the *INS* locus in H3-derived SC-islets (Stage6). (B) Measurement of insulin content per H1-derived SC-islet (n = 4) and primary islet (n = 5). Each datapoint denotes SC-islets from one bioreactor or primary islets from one donor. Statistical significance was tested using a two-tailed unpaired Student’s *t* test with Welch’s correction.

**Figure S3.**
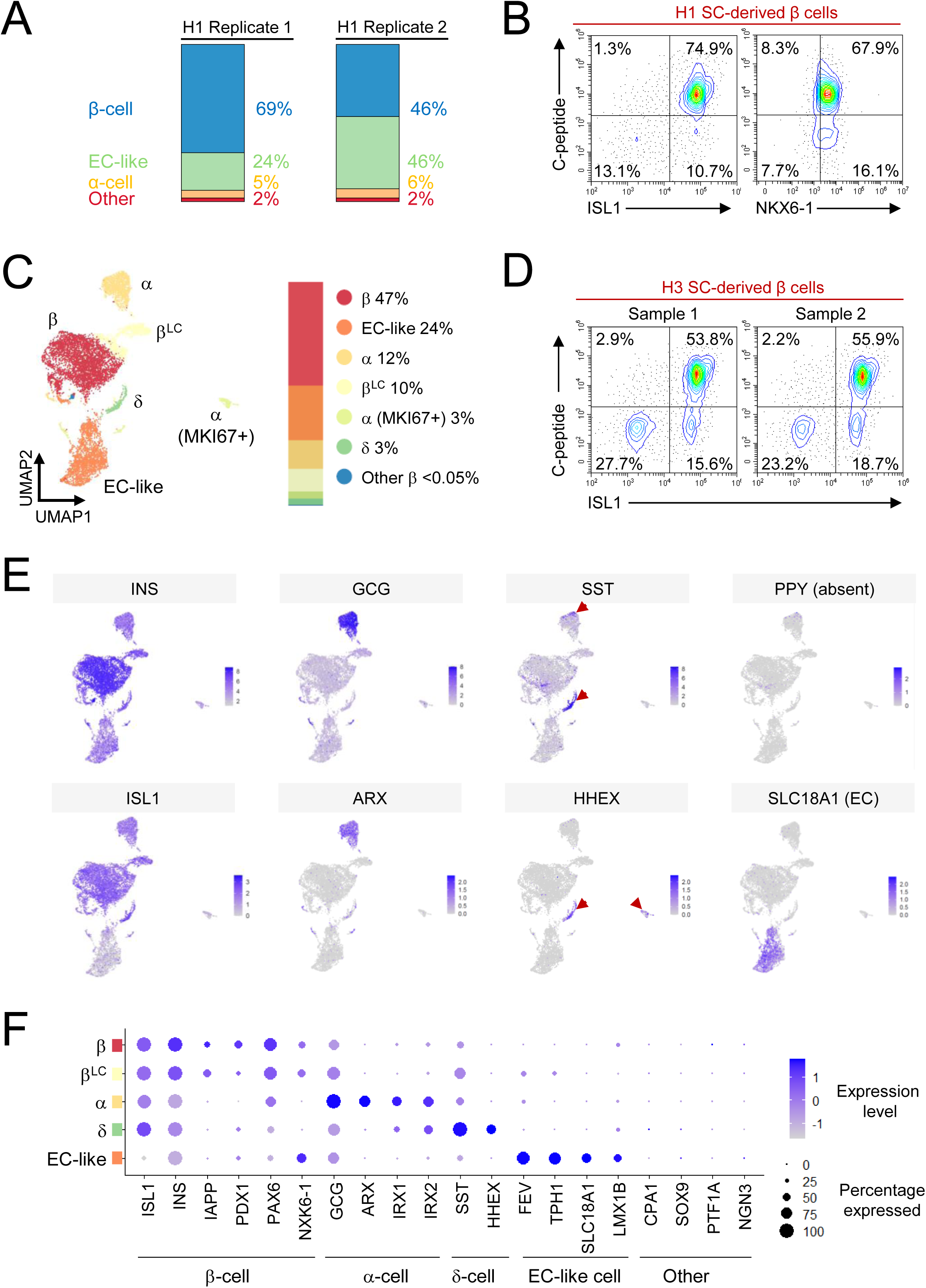
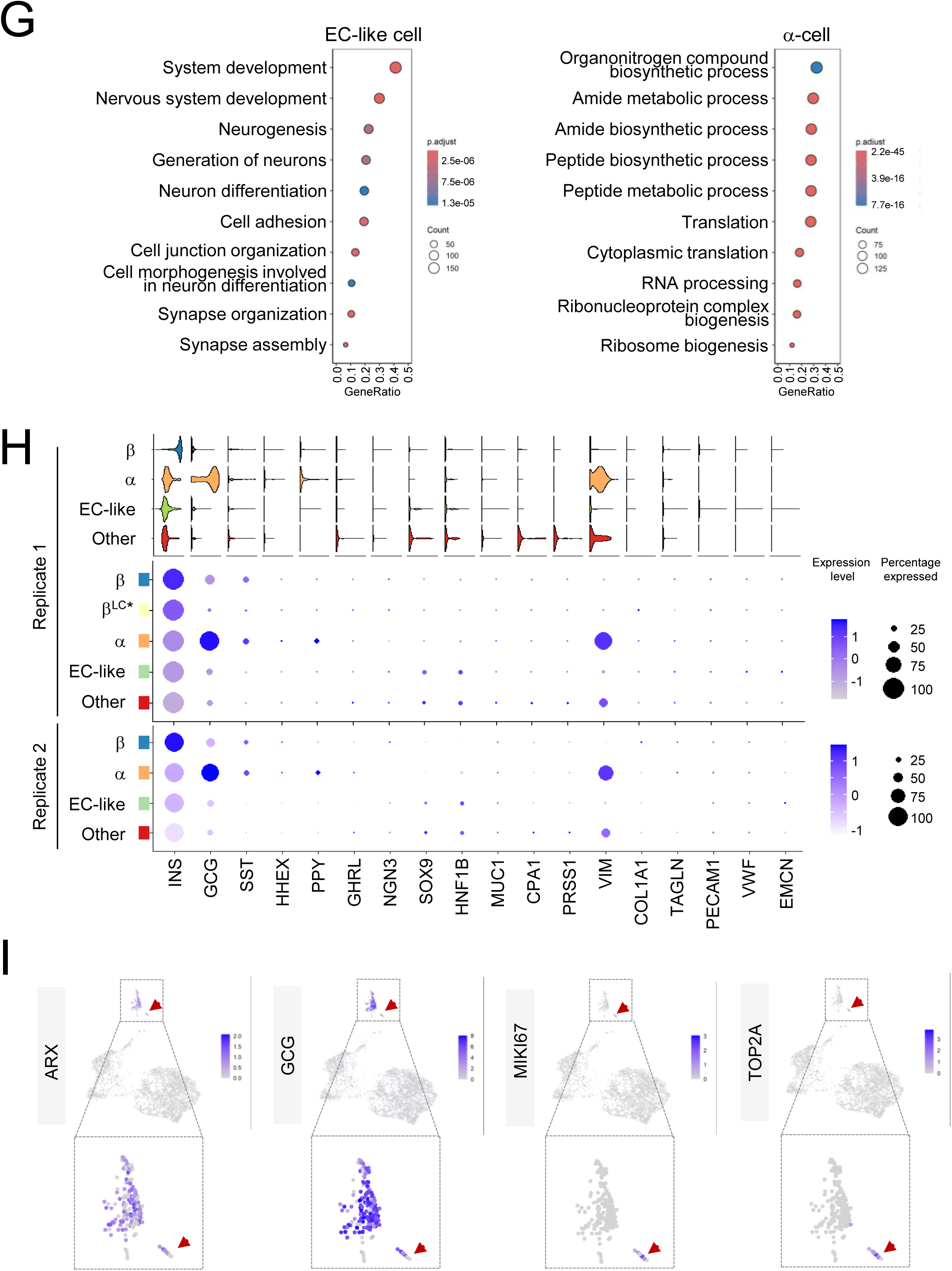
SC-islets lack significant off-target lineage specification by scRNA-seq. (A) percentage of cells in each cluster identified in UMAP projection of scRNA-seq data from two replicates of SC-islets derived from the H1 hESC line at Stage7w2. (B) Flow cytometry contour plots showing the expression of C-peptide, ISL1 and NKX6-1 of SC-islets shown in (A). (C) UMAP visualization of scRNA-seq data from H3 hESC-derived SC-islets at Stage7w2 with percentage of cells in each cluster displayed on the right panel. Clusters were annotated by cell type based on expression of islet hormone genes and established enterochromaffin (EC) cell markers. Please note that β^LC^ cluster with β-cells and exhibit lower total RNA counts, although these cells remained above the filtering threshold for removal. (D) Flow cytometry contour plots showing the expression of C-peptide and ISL1 of SC-islets shown in (C). (E) Feature plots showing expression of key endocrine markers, including INS, GCG, SST, PPY (absent), the pan-endocrine marker ISL1, the α- and δ- cell transcription factors ARX and HHEX, and the EC-cell marker SLC18A1 for H3-derived SC-islets. (F) Dot plot summarizing average expression and the proportion of cells expressing marker genes associated with expected endocrine and off-target lineages across clusters for the H3-derived SC-islets at Stage7w2. (G) Gene Ontology analysis of clusters 1 and 2, corresponding to (left) enterochromaffin (EC)-cell and (right) α-cell populations, respectively, performed using ClusterProfiler R package. (H) (Top panel) Violin plots showing expression of key endocrine hormones and markers across clusters, alongside markers of potential off-target lineages associated with SC-islet differentiation protocols, for H1-derived SC-islet replicate 1; (Middle and lower panels) Dot plots displaying average expression levels and the percentage of cells expressing each marker within clusters, indicating minimal off-target lineage signatures in replicates 1 and 2 of H1-derived SC-islets. (I) scRNA-seq feature plots illustrating expression of selected proliferation and α-cell markers in H1 human embryonic stem cell-derived SC-islets at Stage7w2, with enlarged views of relevant clusters.

**Figure S4.**
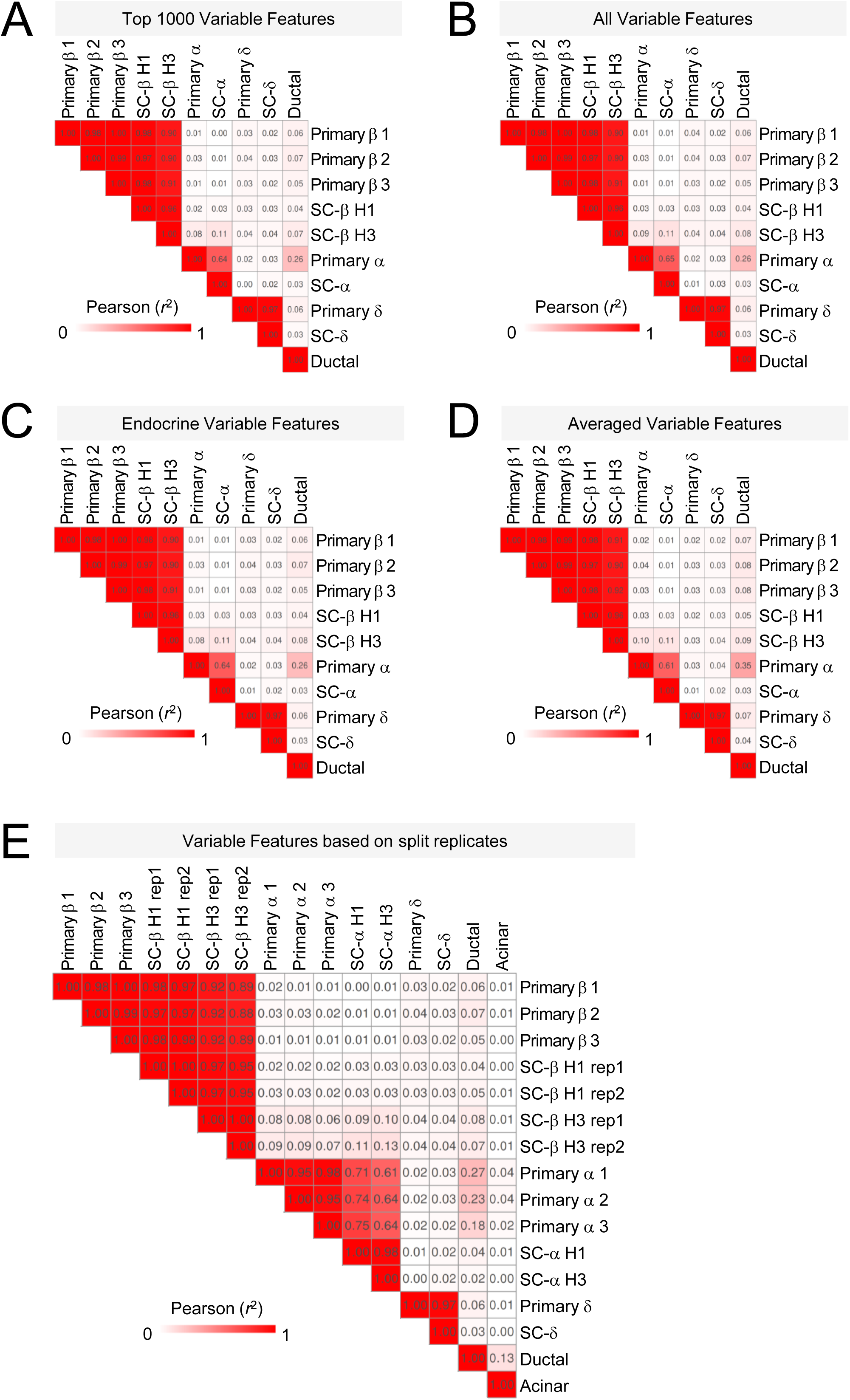
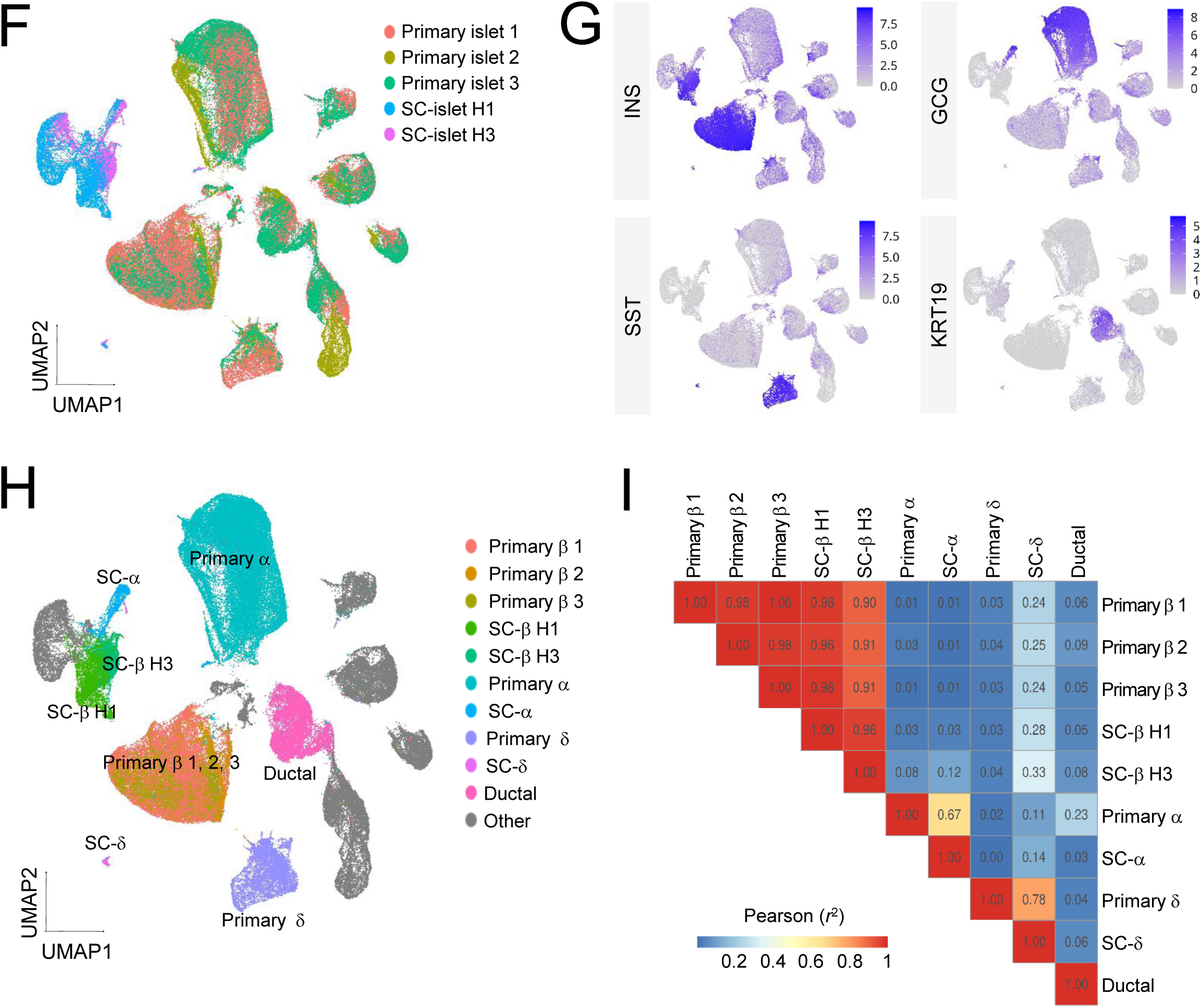
Transcriptional similarity between primary and SC-β cells is reproducible across cell lines and different interrogation methods. (A-E) Heatmaps of the Pearson correlation coefficients of determination (*r*²) calculated by comparing the summarized RNA counts of each cell population‘s (figure 4C) expression of (A) aggregated top 1000 HVFs of all cells in the dataset (all correlation’s BH adjusted p-value < 0.05), (B) all features aggregated (all correlation’s BH adjusted p-value < 0.0001), (C) aggregated top 3000 endocrine (α, β, γ, δ and ε) HVFs (all correlation’s BH adjusted p-value < 0.001), (D) averaged top 3000 HVFs of all cells in the dataset (all correlation’s BH adjusted p-value < 0.0001), (E) aggregated top 3000 HVFs of all cells in the dataset and split into replicates (all correlation’s BH adjusted p-value < 0.01). (F) Combined SCT normalized UMAP visualization of scRNA-seq from 3 publicly available healthy primary islet datasets, and SC islets derived from H1 and H3 hESC lines. Clusters of cells have been annotated by dataset. (G) Feature plots showing expression of key endocrine markers, including INS, GCG, SST, and ductal cell marker KRT19, for (F). (H) Annotated cell types based on dataset (F) and marker expression (G). (I) Heat map of the Pearson correlation coefficients of determination (*r*²) calculated by comparing the aggregated RNA counts of each cell population’s (H) expression of the top 3000 HVFs of all cells in the dataset.

**Figure S5.**
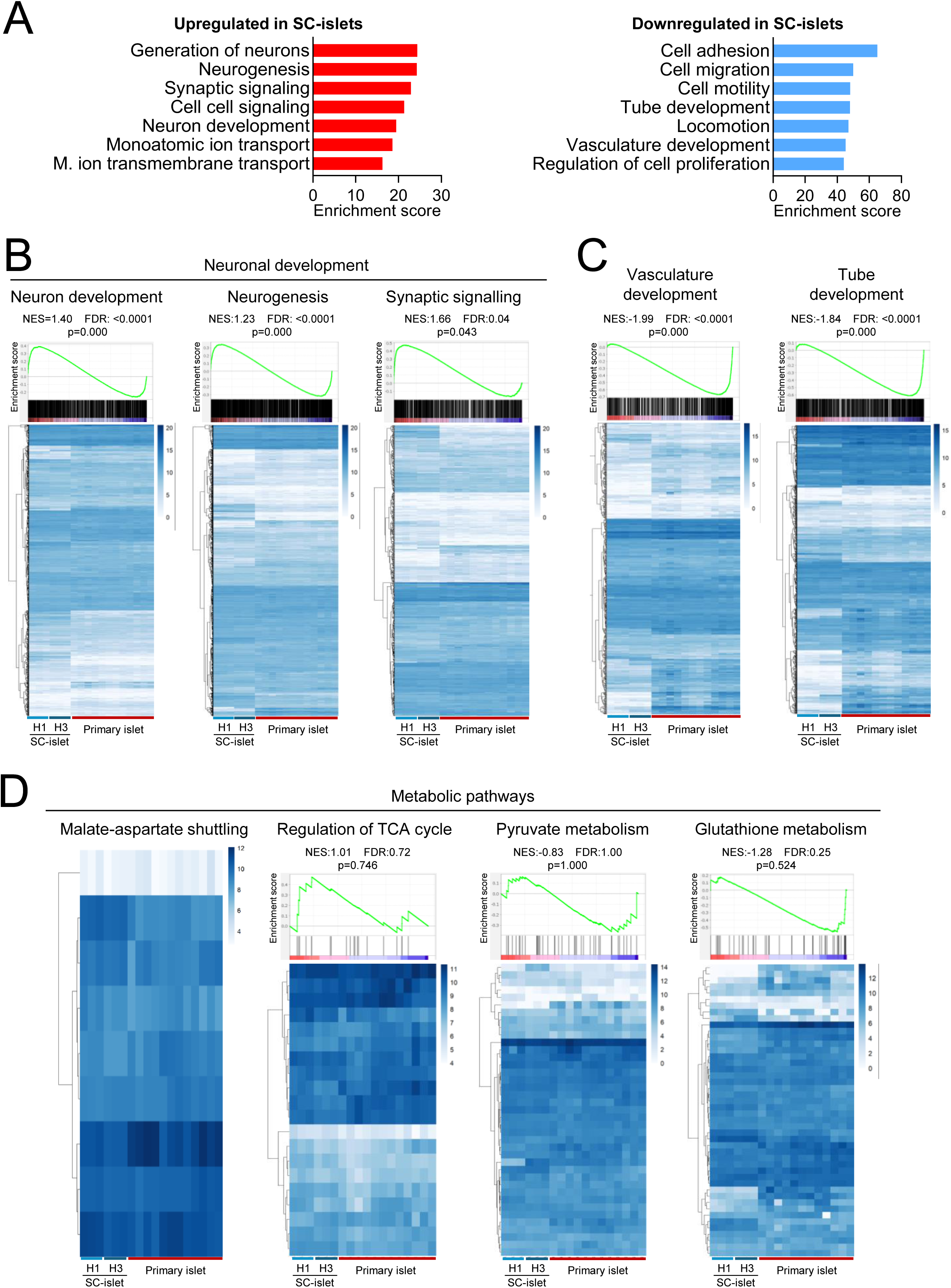
Transcriptional expression of gene pathways associated with β-cell function in SC-islets and donor-derived β cells. (A) Gene Ontology analysis of differentially expressed genes identified by comparing Stage7w2 SC-islets (derived from H1 or H3 hESC lines) with human donor-derived islets (n = 6 SC-islet samples vs n = 12 donor islets). Using DAVID bioinformatics tool (BP) (B-D) (top panels) GSEA analysis for enrichment of indicated gene pathways in SC-islets relative to primary islets and (bottom panels) heatmap showing relative expression of gene sets associated with the indicates gene pathways. Expression values are shown as log2-transformed normalized reads calculated with DeSeq2 in R.

**Figure S6.**
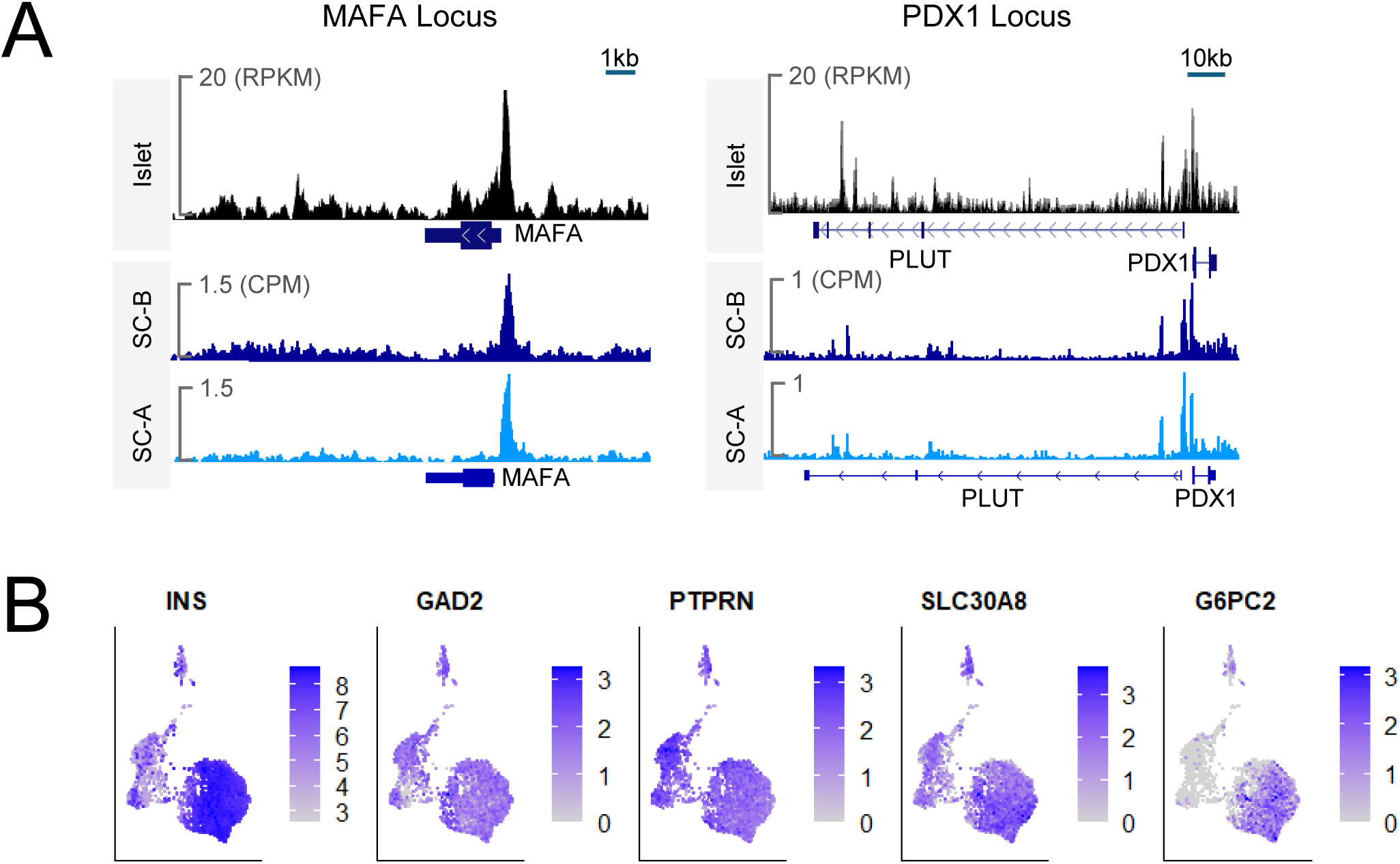
SC-islets appear poised for MAFA expression and express dominant type 1 diabetes antigenic peptides. (A) Genome browser snapshots of ATAC-seq signal at the *MAFA* (left) and *PDX1* (right) loci in human donor–derived islets^67^ and two SC-islet H1-derived Stage7w2 samples. (B) UMAP plots from H1-derived SC-islets scRNA-seq analysis showing expression of indicated genes, including common T1D antigenic peptides.

